# High-recombining genomic regions affect demography inference

**DOI:** 10.1101/2024.02.05.579015

**Authors:** Jun Ishigohoka, Miriam Liedvogel

## Abstract

Inference of population history of non-model species is important in evolutionary and conser- vation biology. Multiple methods of population genomics, including those to infer population history, are based on the ancestral recombination graph (ARG). These methods use observed mutations to model local genealogies changing along chromosomes. Breakpoints at which genealogies change effectively represent the positions of historical recombination events. How- ever, inference of underlying genealogies is difficult in regions with high recombination rate relative to mutation rate. This is because genealogies cover genomic intervals that are too short to accommodate sufficiently many mutations informative of the structure of the un- derlying genealogies. Despite the prevalence of high-recombining genomic regions in some non-model organisms, such as birds, its effect on ARG-based demography inference has not been well studied. Here, we use population genomics simulations to investigate the impact of high-recombining regions on ARG-based demography inference. We demonstrate that inference of effective population size and the time of population split events is systematically affected when high-recombining regions cover wide breadths of the chromosomes. We also show that excluding high-recombining genomic regions can practically mitigate this effect. Finally, we confirm the relevance of our findings in empirical analysis by contrasting demography inferences applied for a bird species, the Eurasian blackcap (*Sylvia atricapilla*), using different parts of the genome with high and low recombination rates. Our results suggest that demography inference using ARG-based methods should be carried out with caution when applied in species whose reference genomes contain long stretches of high-recombining regions.

## Introduction

Population history affects the patterns of genetic variation, and conversely observed genetic variation in genomes allows inference of historical demographic parameters. The increasing availability of genome data of various species at a population level has facilitated development and application of a number of population genomics methods for demography inference (Excoffier et al., 2013; Gutenkunst et al., 2009; Harris & Nielsen, 2013; Li & Durbin, 2011; Liu & Fu, 2020; Schiffels & Durbin, 2014; Terhorst et al., 2017). These approaches are typically first applied to human data to understand the population history of our own species and for validation of the new methods (Excoffier et al., 2013; Gutenkunst et al., 2009; Harris & Nielsen, 2013; Li & Durbin, 2011; Liu & Fu, 2020; Schiffels & Durbin, 2014; Terhorst et al., 2017), but thereafter adopted to other species including domesticated and wild organisms to answer evolutionary questions (Alonso-Blanco et al., 2016; Groenen et al., 2012; Lanier et al., 2015; Liu et al., 2014; Nadachowska-Brzyska et al., 2016) and to assist conservation efforts (Dussex et al., 2021; Hohenlohe et al., 2021; Li et al., 2014; Pacheco et al., 2022). Despite the wide application of demography inference methods in non-model organisms, their performance outside the parameter space of humans has not been well evaluated.

Some methods for demography inference are based on the ancestral recombination graph (ARG) (Li & Durbin, 2011; Schiffels & Durbin, 2014; Speidel et al., 2019; Terhorst et al., 2017). The ARG is a structure that describes the full ancestries of sampled genomes along recombining chromosomes (Griffiths & Marjoram, 1997). It essentially consists of a series of marginal genealogical trees changing in the topology and branch lengths along the chromosome, and their breakpoints effectively represent historical recombinations contributing to the sampled genomes (Fig. 1). The full ARG provides rich information on the population history (i.e. all coalescence and recombination events through time and mutations mapped on branches), making ARG-based methods a powerful population genomics approach to study evolutionary processes (Hubisz et al., 2020; Schaefer et al., 2021; Speidel et al., 2019; Stern et al., 2019; Wohns et al., 2022). In practice, however, ARG-based methods depend on inference of the ARG (Ignatieva et al., 2021; Kelleher et al., 2019; Mirzaei & Wu, 2017; Rasmussen et al., 2014; Speidel et al., 2019; Wohns et al., 2022), or representations of underlying genealogies (Li & Durbin, 2011; Schiffels & Durbin, 2014; Terhorst et al., 2017), which in turn relies on observed mutations. Importantly, the presence of mutations representing an ARG branch depends on recombination and mutation rates. If an ancestral haplotype breaks by a recombination before accommodating mutations, the corresponding branch on the ARG is not represented by any mutations (Fig. 1B) (Hayman et al., 2023; Shipilina et al., 2023). Therefore, high recombination rates (relative to the mutation rate) makes it difficult to accurately infer the underlying ARG, limiting the performance of the ARG-based approach (Sellinger et al., 2020, 2021; Terhorst et al., 2017).

**Figure 1:**
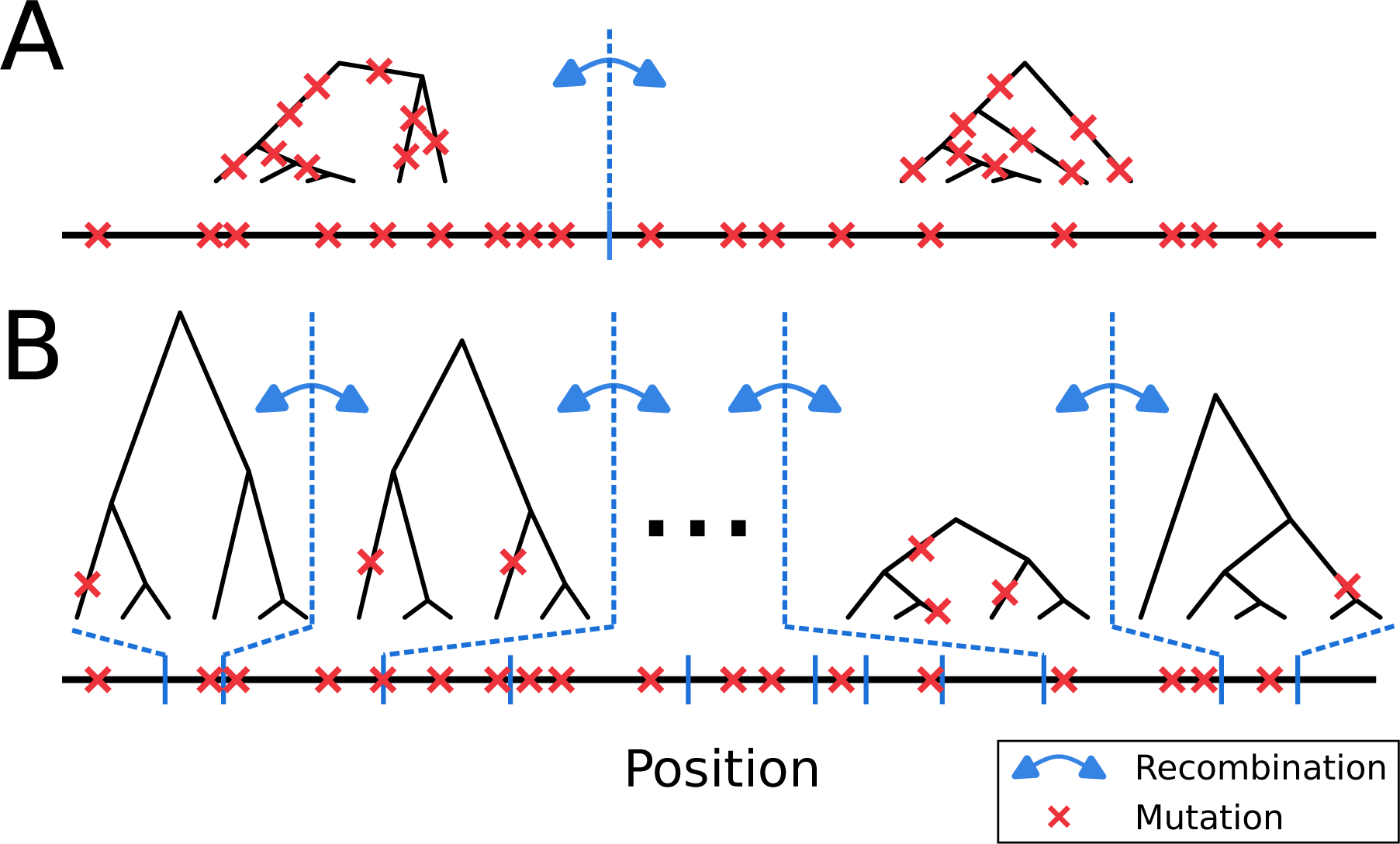
**The presence of mutations representing ARG branches depends on recombination rate**. **A**. When recombination rate is moderately low, branches of ARG are represented by mutations. This allows inference of the underlying ARG based on observed mutations. **B**. When recombination rate is high, many branches of ARG are not represented by any mutations. We ask whether this affects ARG-based demography inference.

The impact of high recombination rate on ARG-based demography inference is presumably negligible in humans (Li & Durbin, 2011; Terhorst et al., 2017) where recombination rate is low except for narrow recombination hotspots (Myers et al., 2010; Stevison et al., 2016). However, this type of recombination landscape is not universal to all organisms (Auton et al., 2013; Baker et al., 2017; Lam & Keeney, 2015; Singhal et al., 2015), including species with ecological and evolutionary relevance or of conservation concern. The difference in recombination landscapes can be partially attributed to the presence and absence of PRDM9, a transcription factor that determines the genomic position of recombination hotspots. PRDM9 introduces histone modifications to recruit the molecular machinery initiating DNA double-strand breaks (DSBs), which is required for meiotic recombination (Baker et al., 2015; Baudat et al., 2010; Paigen & Petkov, 2018). PRDM9 has been lost independently at least thirteen times in vertebrates (Cavassim et al., 2022), which shifted the recombination hotspots from rapidly evolving PRDM9 motifs (Baker et al., 2015; Myers et al., 2010; Oliver et al., 2009) to genome features such as transcription start sites and CpG-islands (Auton et al., 2013; Baker et al., 2017; Kawakami et al., 2017; Paigen & Petkov, 2018; Singhal et al., 2015). Hotspots of PRDM9-independent recombination in birds (Bascón-Cardozo et al., 2024; Kawakami et al., 2017; Singhal et al., 2015), dogs (Auton et al., 2013) and percomorph fish (Baker et al., 2017) appear to be wider than PRDM9-dependent hotspots in primates (Durbin et al., 2010; Myers et al., 2010; Stevison et al., 2016). On top of the recombination landscape, the average recombination rate is highly variable between chromosomes and species (Stapley et al., 2017). These differences in meiotic recombination could potentially impact modelling of the local ARGs in non-human species.

In this study, we ask how recombination landscapes affect ARG-based demography inference. To this end, we simulate genome data under a simple demographic history with various recombination maps, and evaluate the accuracy of demography inference by different ARG-based methods. Specifically, we focus on two ARG-based methods, MSMC2 (Malaspinas et al., 2016; Wang et al., 2020) and Relate (Speidel et al., 2019), differing in the way the ARG is modelled. While Relate infers a series of marginal genealogies along the genome with their topology and branch lengths collectively representing the full ARG of the sample, MSMC2 models the distribution of the coalescence times between pairs of sampled sequences along the genome based on the sequentially Markovian coalescent (SMC, McVean & Cardin (2005)). To demonstrate the relevance of our findings based on simulations, we translate our findings to empirical data of a non-model organism with wide high-recombining genomic regions. To this end, we use whole-genome resequencing (WGR) data and fine-scale recombination maps of a songbird species, the Eurasian blackcap (*Sylvia atricapilla*), and contrast ARG-based demography inferences using genomic regions differing in recombination rates.

## Results

### Simulations of different recombination maps

To investigate the effect of the recombination landscape on ARG-based demography inference, we used msprime (Kelleher et al., 2018) to simulate a simple demographic history (Fig. 2A) with five different recombination maps (Fig. 2B). In all simulations, three subpopulations (pop1, pop2, and pop3) split from a constant-sized ancestral population of 1 million diploids 10,000 generations before the present, after which they followed different trajectories of effective population size (constant (pop1), exponential increase (pop2), and exponential decrease (pop3)). Under this demography model, we simulated 16 Mb-long chromosomes (Fig. 2B) with a constant mutation rate and heterogeneous recombination rate along the chromosome (color-coded from orange (no high-recombining regions) to dark red (extreme high-recombining regions)). Specifically, the 10 Mb stretch in the middle of the chromosome (“middle”) had a recombination rate one-tenth the mutation rate in all five scenarios, while the recombination rate increased in a stepwise manner at the 3 Mb ends of the chromosome. To test whether masking high-recombining regions improves ARG-based demography inference, we applied masks accounting for the length of chromosomes in two ways: masking a total of 6 Mb within the central part of the chromosome without elevated recombination rate (“control”, Fig. 2C top) or masking the 3 Mb ends of the chromosome covering the entire high-recombining regions (Fig. 2C bottom). Mean recombination-mutation ratios of the five scenarios for the control condition (after applying the masks) were 0.1, 0.25, 1, 4, and 10 (Fig. 2B, C)). We inferred the demography with two ARG-based methods, MSMC2 and Relate. We also inferred demography using Stairway plot 2, an SFS-based method expected to be unaffected by elevated local recombination rate. In summary, we tested the performance of demography inference in grids of five scenarios (differing in recombination maps), three demography inference methods (two ARG-based and one SFS-based), and two conditions of masking (masking high-recombining and background regions).

**Figure 2:**
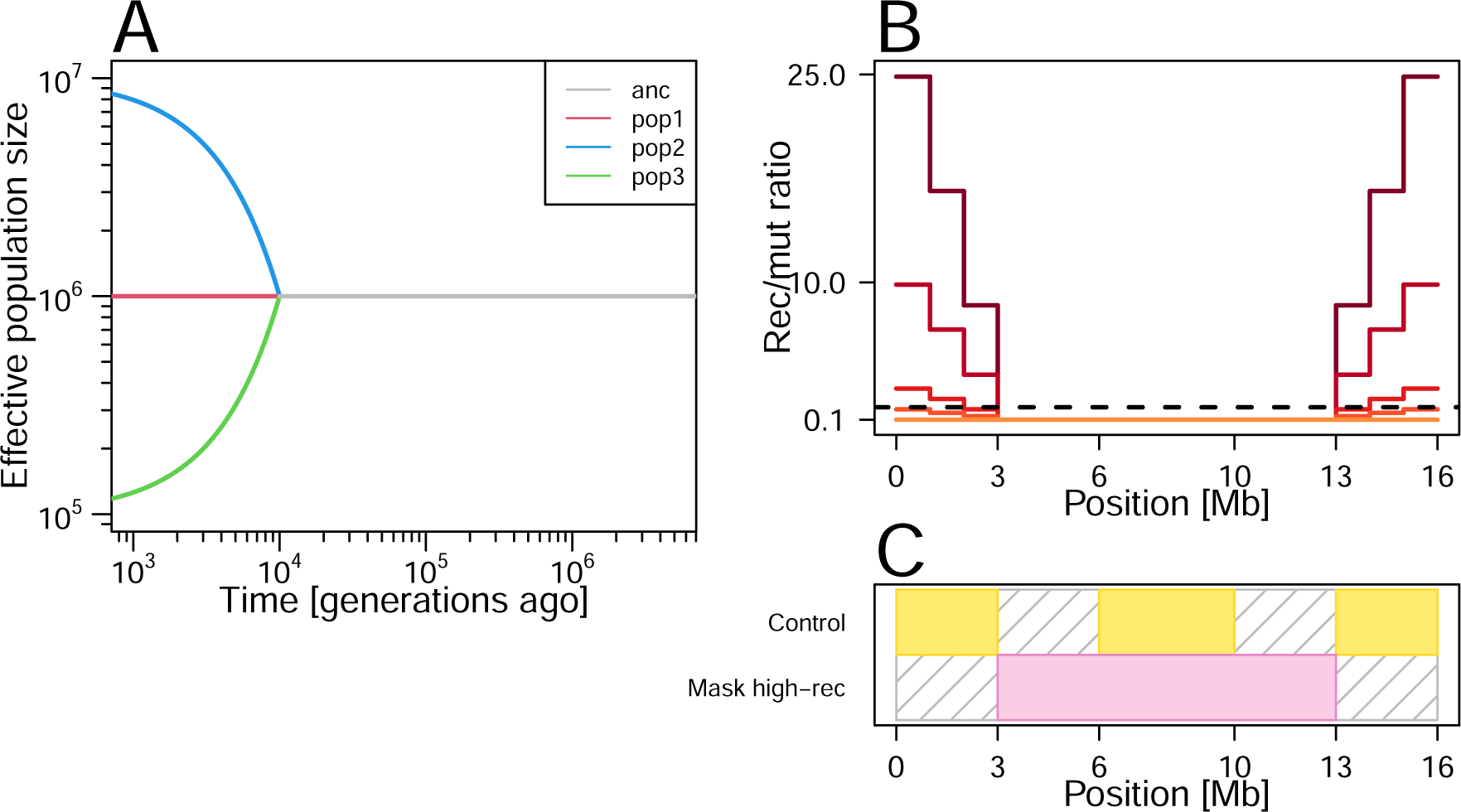
Design of our simulation study. A. Simulated demographic history. We simulated three populations (pop1, pop2, pop3) that split from one ancestral population simultaneously 10,000 generations before the present. After the population split, these three populations follow distinct trajectories of effective population size. **B.** Recombination maps used in our simulations for a hypothetical chromosome. We considered five scenarios with different recombination maps that are depicted by five solid color-coded lines. In all scenarios, recombination rate (Y axis) is 1/10 of the mutation rate in the middle of the chromosome. Recombination rate increases in a stepwise manner (scenarios with different levels of increase are color-coded from orange to dark red) towards the ends of the chromosome. The horizontal black dotted line indicates the recombination-to-mutation ratio of 1. **C.** Two settings of demography inference masking different parts of the chromosomes (yellow blocks depict genomic regions used for demography inference. Gray shades depict regions masked from demography inference). We asked whether the presence of high-recombining regions (i.e. towards the chromosome ends in the simulated scenario illustrated in C) in the genome affects demography inference and whether masking them improves demography inference. We applied methods of demography inference on the simulated data using different parts of the chromosome: either including the high-recombining regions (“control”, top) or masking the 6 Mb of high-recombining regions (bottom). To control for the total sequence length used in the inferences, we applied a total of 6 Mb masks outside the high-recombining regions in the first setting (top).

### Effect of wide high-recombining regions on inference of effective population size

We first investigated the effect of high-recombining regions on the inference of historical effective population size under the control condition, in which high-recombining regions were not masked (Fig. 3A-C). Inference by MSMC2 was too noisy in the recent past to inspect the timing of the split event based on inferred effective population size especially in scenarios with mild or no elevation of recombination rates at the chromosomal ends (Fig. 3, left column. Results of all five recombination landscapes are in Sup. Fig. 1). This noisiness in MSMC2 inference is unlikely to be the effect of recombination rate, as the inference was noisy even for the scenario without a high-recombining region (Fig. 3A, left column), but rather results from insufficient data (eight haploids of ten chromosomes of 16 Mb) and the simulated split time being too recent compared to the effective population size. When the mean recombination rate was lower than or equal to the mutation rate, MSMC2 inferred effective population size accurately in the deep past (Fig. 3A, B, left column). However, when the mean recombination rate was greater than the mutation rate, the inference of effective population size by MSMC2 was systematically deviated in the deep past with a characteristic wave-shaped pattern in the skyline plot (Fig. 3C, left column). Inference by Relate, which exploited a total of 300 haploids from the three subpopulations, was less noisy compared to MSMC2 even in the recent past (Fig. 3, middle column. Results of all five recombination landscapes are in Sup. Fig. 1). In the case without high-recombining regions (Fig. 3B, middle column), the apparent split time as well as the post-split trajectories of the effective population sizes were accurately inferred, albeit slightly smaller effective population size before the split event was estimated. However, effective population sizes of pop1 (red, constant size) and pop2 (blue, exponential growth) were systematically underestimated after the population split as the mean recombination rate increased. In contrast to the two ARG-based methods, demography inference by SFS- based Stairway plot 2 was robust to the presence of high-recombining regions (Fig. 3, right column).

**Figure 3:**
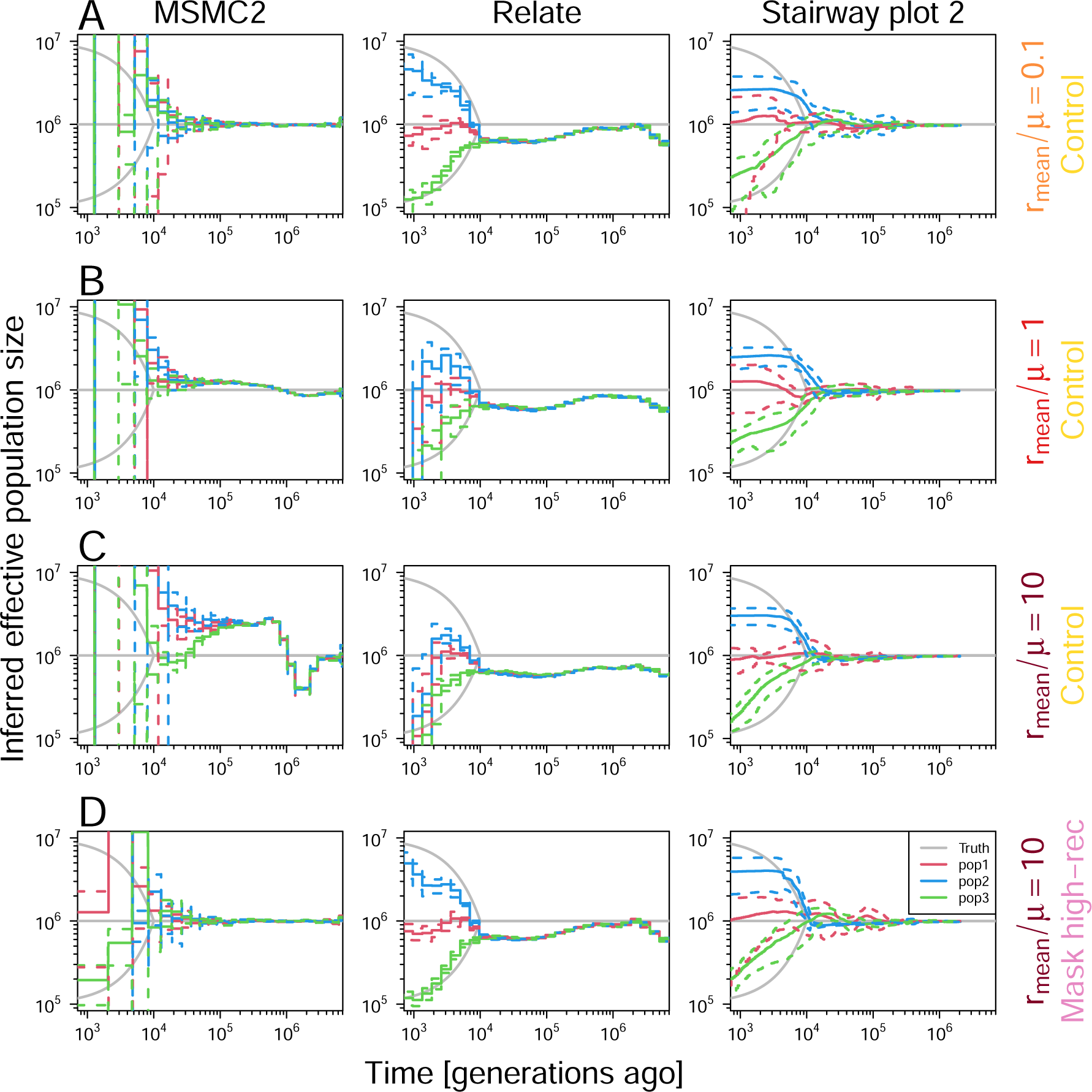
Inference of historical effective population size. The inferences by MSMC2 (left) and Relate (middle) without masking high-recombining regions (**A-C**) show that ARG-based methods are biased in the presence of high-recombining regions. Removing the high-recombining regions eliminated the bias (**D**). The results for Stairwayplot 2 (right) confirm the expectation that the SFS-based method is less affected by the presence of high-recombining regions. In each panel, gray lines depict the simulated truths (as in Fig. 2A), solid and dashed colored lines depict the mean and mean *±* SD of the inferences (see Materials and Methods for details on how replicates were treated).

To investigate whether the deviations in the demography inferences by ARG-based methods are due to errors in inferred local ARGs within the high-recombining regions or global errors throughout the entire chromosome, we compared coalescence time metrics in inferred (representation of) ARGs with the simulated truth. For MSMC2, we compared discretized time to the most recent common ancestor (TMRCA) between a pair of haplotypes along chromosomes with the true coalescence times. For Relate, we compared TMRCA of the entire ARG of 300 sequences along chromosomes between the inference and the truth. In both ARG-based methods, correlation between the inference and the truth was reduced specifically in the high-recombining regions (Sup. Figs. 3, 4). Furthermore, masking high-recombining regions improved demography inference in both methods (Fig. 3D). Our findings indicate that ARG-based demography inference is affected by localized errors within the high-recombining regions.

### Effect of wide high-recombining regions on the inference of population splits

Next, we investigated the effect of high-recombining regions on ARG-based inference of population split events under the control condition (without masking high-recombining regions) between pairs of populations by inferring historical relative cross-coalescence rate (rCCR) by MSMC2 and Relate (Fig. 4A-C. Results of all five recombination landscapes are illustrated in Sup. Fig. 2). rCCR is a metric informative of split events and gene flow between a pair of populations: it increases from 0 to 1 backwards in time (i.e. it drops from 1 to 0 forward in time) at the population split event, and the rate of this change in respect to time essentially depicts how fast the split occurred (Schiffels & Durbin, 2014).

**Figure 4:**
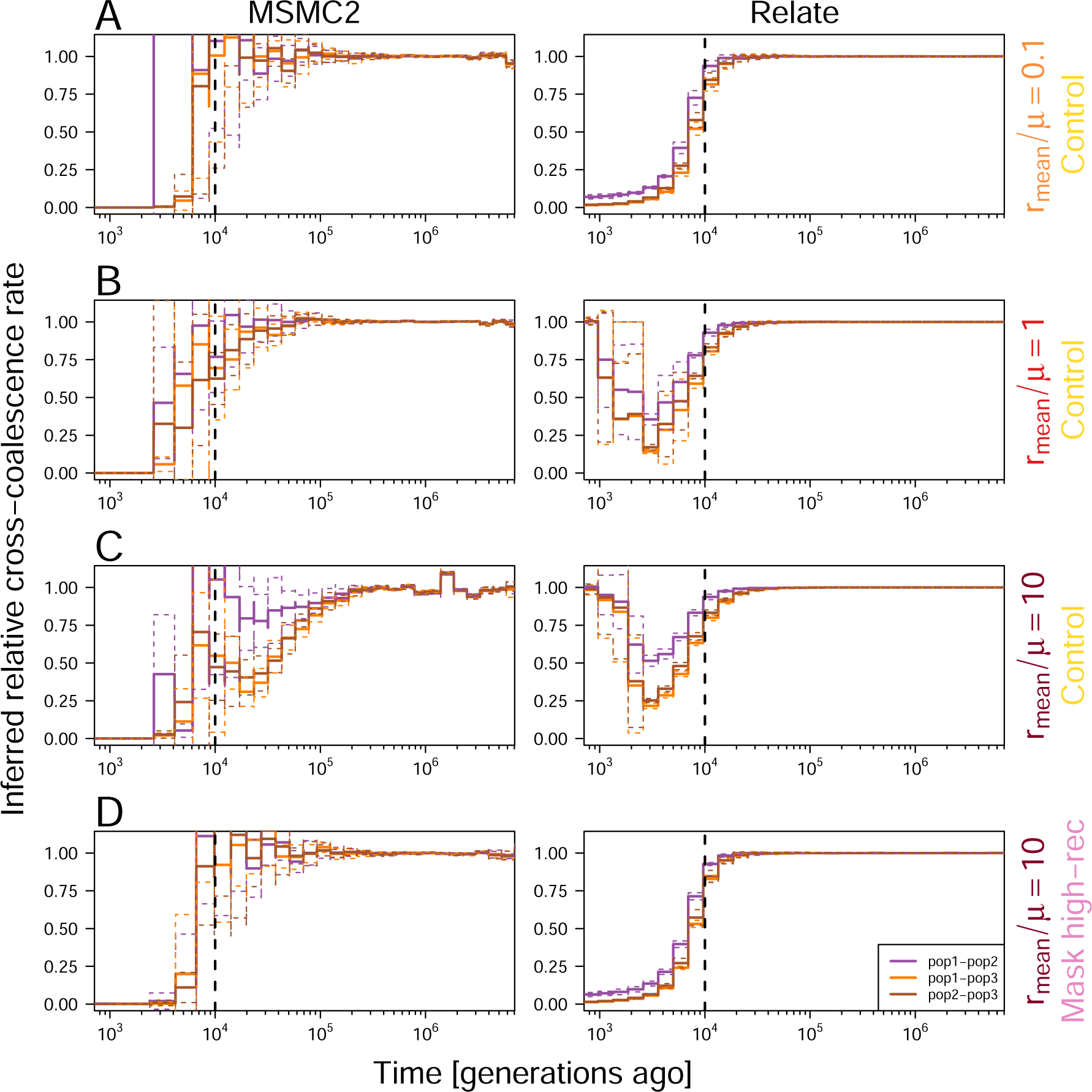
Inference of population split events. Vertical dotted lines depict the true split time. Colored lines depict inferred rCCR for pairs of populations. Three colors indicate three pairs of populations. **MSMC2** (left column). Solid lines depict mean of inferences of the ten down-samples, and dotted lines depict mean *±* SD. The results show that the presence of high-recombining regions biases the inference of population splits, and removing the high-recombining regions reduces this bias. **Relate** (right column). Solid lines depict mean of inferences of the ten replicates, and dotted lines depict mean *±* SD. The results show that the presence of high-recombining regions biases the inference of population splits, and removing the high-recombining regions reduces this bias.

Inferred rCCR based on MSMC2 started to drop (forward in time) earlier than the true split time (Fig. 4). This was especially true for subpopulation pairs involving pop3 (exponential reduction in effective population size) as the mean recombination rate increased. This pattern is consistent with the older apparent split based on effective population size inferred by MSMC2 with high-recombining regions (Fig. 3C, left column). In inference by Relate, rCCR decreased (forward in time) at the true split time, but increased again towards the present time in scenarios with high-recombining regions (Fig. 4). Importantly, we were able to mitigate these effects by masking the high-recombining regions (Fig. 4D). To summarize, our simulation study illustrates that the presence of high-recombining regions affects ARG-based inference of both historical effective population size and population split time. We could show that these effects can be mitigated by masking high-recombining regions of the genome in the analysis.

### Effect of a narrow high-recombining region on demography inference

To further explore how different levels of heterogeneity in the local recombination rate along chromosomes might affect ARG-based demography inference, we simulated the same demo-graphic history with two additional sets of recombination maps. Both sets consisted of five recombination maps (Sup. Fig. 5): the first set with uniform recombination rates along the chromosome (“Uniform”. 0.1, 0.25, 1, 4 and 10 times the mutation rate); and the second with a single narrow region of high recombination rate Fig. 2B (“Narrow high-rec.”. Ratios between mean recombination rate and mutation rate of 0.1, 0.25, 1, 4, and 10 as in Fig. 2 but with a narrower region of high recombination rate). For the uniform scenario, inference by both MSMC2 and Relate were affected by recombination rate similarly to the stepwise recombination maps as the recombination rate increased (Sup. Fig. 6). For the narrow high-rec. scenario, inference by MSMC2 was slightly affected when the mean recombination rate was greater than the mutation rate (Sup. Fig. 7). In contrast, inference by Relate was robust to the presence of the narrow high-recombining region. These results indicate that the ARG-based methods are affected by a wide coverage of regions with recombination rate greater than mutation rate. Additionally, the ARG-based methods differ in their robustness to the presence of high-recombining regions. This may represent an inherent difference of the robustness between MSMC2 and Relate, but it also could reflect the different sample sizes used as the input owing to different scalability between the two methods: in our study we used 150 diploids (300 haploid genomes) for inference by Relate, while only eight haploid genomes per population were used for MSMC2.

### Application to empirical data

Birds lack PRDM9 and their recombination is characterized by higher rates and wider hotspots compared to PRDM9-dependent recombination in primates (see Introduction). Additionally, per-generation mutation rate is lower in many birds than in humans (Bergeron et al., 2023). These factors are manifested as wider breadth of genomic regions with high recombination- mutation ratio, which could bias ARG-based methods for demography inference as shown in our simulation study. To address the relevance of our findings to empirical applications, we revisited genome data of the Eurasian blackcap (*Sylvia atricapilla*, “blackcap” here after) as a representative of birds. The data consist of whole-genome resequencing of 179 individuals of ten populations across the species’ distribution range (Delmore et al., 2020; Ishigohoka et al., 2023), covering variation in population-typical phenotypes of seasonal migration. To investigate the effect of recombination rate on ARG-based methods of demography inference with the blackcap dataset, we split the blackcap genome into low- and high- recombining halves, based on local recombination rates characterized in a previous study (Bascón-Cardozo et al., 2024) (see Materials and Methods for details). We performed demography inference (effective population size and rCCR) with MSMC2 and Relate for each half separately.

Inference by MSMC2 showed apparent effects of high-recombining regions consistent with our simulation. Historical effective population size inferred using the high-recombining half had characteristic wave-shaped trajectory in the deep past of the skyline plot (Fig. 5B) compared to that using the low-recombining half (Fig. 5A). The apparent split time between populations based on effective population size was older using the high-recombining half (Fig. 5B) than using the low-recombining half (Fig. 5A). In line with this, direct comparison of inferred rCCR for pairs of blackcap populations between the high- and low-recombining halves revealed inference of systematically older split time using the high-recombining half than the low-recombining half (Fig. 5C, D). These differences are consistent with our simulation study (Figs. 3, 4), indicating that the effect of high-recombining regions on inference by MSMC2 is relevant for empirical analysis. In contrast, Relate was robust to the difference in recombination rate between the two conditions (Sup. Fig. 8). This difference between MSMC2 and Relate suggests different levels of robustness of ARG-based methods to the presence of high-recombining regions, which is in line with our simulation study.

**Figure 5:**
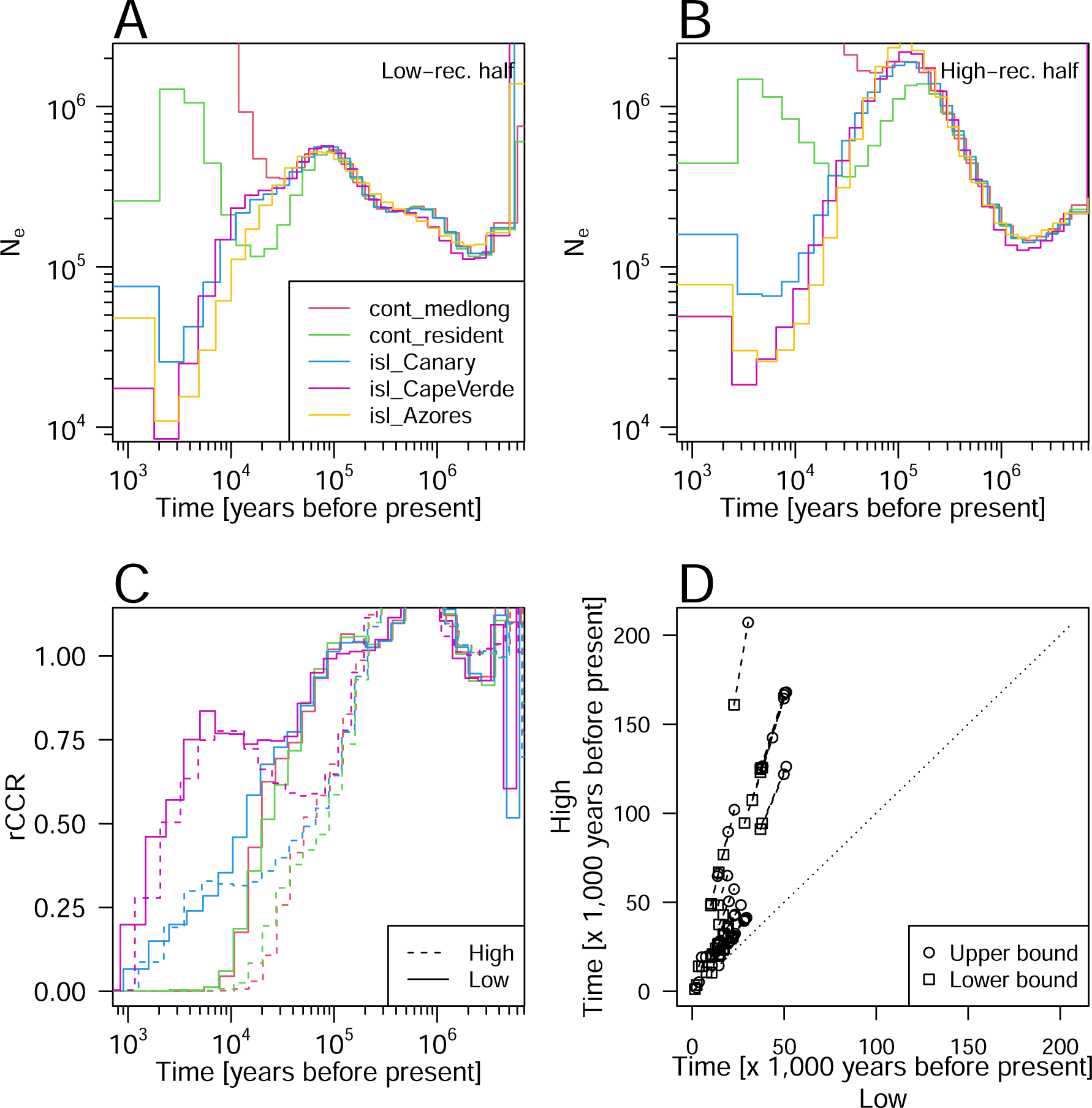
**High-recombining regions can affect demography inference in empirical analysis. A, B**. Inference of historical effective population size by MSMC2. Results for five exemplified blackcap populations are shown using the lower (**A**) or the higher (**B**) half of the genome based on local recombination rates. **C**. Inference of relative cross-coalescence rate (rCCR) with MSMC2 between Azores population and each of all other four populations in **A** and **B** using the lower (solid lines) and the higher (dotted lines) half of the genome based on local recombination rates. **D**. Comparison of split times inferred by MSMC2 using the lower higher halves of the genome based on recombination rates. Segments represent inference between 45 pairs of 10 populations. Two ends of a segment represent the lower and upper boundaries of two consecutive discretized epochs between which rCCR crosses the threshold of 0.5.

## Discussion

Our results suggest that demography inference using ARG-based methods should be carried out with caution in organisms that are likely to harbor recombination landscapes distinct from humans, for which these methods were initially developed. In many animals with functional PRDM9, including humans, recombination events are concentrated in narrow recombination hotspots (Myers et al., 2010; Stevison et al., 2016). Thus, we expect that ARG-based methods will be robust. Other species have high-recombining regions more widely distributed around genomic features along the genome (Baker et al., 2017; Kawakami et al., 2017; Singhal et al., 2015) and thus ARG-based methods can be more susceptible to the effect of high-recombining regions. Using simulations we demonstrated that masking high-recombining regions improves the ARG-based demography inference in such cases. In practice, however, this raises another question of how to define regions to mask, which can be challenging due to multiple factors. First, defining a threshold value of the recombination rate using inferred recombination maps may be problematic. This is because methods for inference of fine-scale recombination maps can be inaccurate, especially when the recombination rate is higher than the mutation rate (Raynaud et al., 2023; Spence & Song, 2019). Second, long-enough contiguous chromosomal segments are essential in ARG-based methods (Sellinger et al., 2021), hence masking every high-recombining region can be problematic as it might split the genome into pieces too small for ARG-based methods to be applied. An additional factor to take into account is variation among chromosomes. For example, in multiple taxa, the chromosome lengths can substantially vary, and the recombination rate is negatively correlated with the chromosome length (Bascón-Cardozo et al., 2024; Kawakami et al., 2014; Martin et al., 2019; Singhal et al., 2015), potentially leading to different applicability of ARG-based methods among chromosomes. Finally, additional masks may be necessary for demography inference if large blocks that do not represent neutral evolution exist in the genome. For example, large polymorphic inversions under long-term balancing selection (Giraldo-Deck et al., 2022; Hager et al., 2022; Harringmeyer & Hoekstra, 2022; Kim et al., 2017; Knief et al., 2016, 2017; Küpper et al., 2015; Lamichhaney et al., 2015; Mérot et al., 2021) may be excluded, which may leave little data for inference in species with small genomes. We suggest to run simulations tailored to the species under study to assess whether ARG-based methods can be used with some confidence. This is especially necessary in species without functional PRDM9, with broad high-recombining regions, high genome-wide mean recombination rates, small genomes, highly heterogeneous chromosomes, and large structural variations.

An SFS-based method for demography inference, Stairway plot 2, performed well without high-recombining regions and even better without high-recombining regions in our simulation study. We propose that this accuracy under the presence of high-recombining regions can represent a general characteristic of SFS-based methods that they benefit from high-recombining regions, from which ARG-based methods suffer. The problem of high-recombining regions for ARG-based methods is the fact that branches of genealogies are not represented by mutations (Hayman et al., 2023). In other words from the perspective of mutations, ARG-based methods suffer from independence of mutations in a local genomic window. This independence of mutations, however, is the assumption to compute SFS (Gutenkunst et al., 2009), allowing SFS-based methods to perform accurately with high-recombining regions. Localized errors in the inference of (representation of) genealogies within high-recombining regions in our study indicate that the issue of high-recombining regions in ARG-based methods is not specific to demography inference but can be critical in other applications, including inference of selection (Hejase et al., 2020; Speidel et al., 2019; Stern et al., 2019). In regions with high recombination rates, inferred genealogies may be too inaccurate to perform ARG-based selection tests, while SFS-based methods may be used on the local variation data (Fay & Wu, 2000; Tajima, 1989). Combining ARG- and SFS-based approaches, giving them complementary weights according to the local recombination rate, may make the most of the variation data in demography inference.

In this study, we demonstrated that the recombination landscape can influence ARG-based approaches of population genomics. Although the true ARGs should have rich information on the population history and evolutionary processes, the effects of errors in inferred local ARGs within regions of elevated recombination rate are, in some cases, not negligible. Our findings are likely relevant not only to birds but likely in a wide range of species, because PRDM9 has been lost at least thirteen times independently in vertebrates (five clades of ray-finned fish, four clades of amphibians, a clade of lizards, the entire clade of birds and crocodiles, and two clades of mammals (dogs and platypus) (Cavassim et al., 2022)). In addition to the recombination rate, other factors, such as the genomic landscape and spectra of mutation (Jiang et al., 2021; Monroe et al., 2022; Sasani et al., 2022; Wu et al., 2020), local effective population size (reflecting selection: Nielsen, 2005; Burri, 2017; Ellegren & Galtier, 2016), and effective migration rate (reflecting barriers to gene flow: Westram et al., 2022) are distributed non-uniformly along the genome, and they may similarly affect population genomics summary statistics and inferences. Novel approaches jointly modelling heterogeneity of some of these factors are emerging (Barroso & Dutheil, 2023; Korfmann et al., 2023; Laetsch et al., 2023). Nonetheless, we highlight that evaluating the performance and limitation of population genomics methods under non-canonical parameter space relevant in individual cases is necessary to draw meaningful interpretations.

## Materials and Methods

### Simulation study

#### Simulation

To investigate the effect of high-recombining regions on demography inference, we simulated ARGs and mutations with msprime version 1.2.0 (Baumdicker et al., 2022) under the standard neutral coalescent with recombination (Hudson, 1983). The demography model consisted of an ancestral population of 1,000,000 diploids splitting into three populations (pop1, pop2, and pop3) at 10,000 generations before the present time. The population size of pop1 was constant at 10,000, and exponential increase and decrease of 10 folds over 10,000 generations were introduced in pop2 and pop3 after the split event. The mutation rate was set to 4.6 *×* 10*^−^*^9^ per generation per site.

We prepared three sets of recombination maps, each of which consists of five scenarios. The first set (“stepwise”) was 16 Mb long, and the recombination rate was set to one tenth the mutation rate (4.6 *×* 10*^−^*^10^) at the central 10 Mb, with a step-wise increase in recombination rate at 3 Mb ends of the chromosome (Fig. 2B), such that the mean recombination rate (after masking 6 Mb of the middle) were 0.1, 0.25, 1, 4, and 10 times the mutation rate. The second set (“narrow high-rec.”) was 11 Mb long, and the recombination rate was set to 4.6 *×* 10*^−^*^10^ throughout the chromosome, except a 1 Mb segment in the middle, where recombination rate was elevated such that the mean recombination rate were 0.1, 0.25, 1, 4, and 10 times the mutation rate Sup. Fig. 5A. The third set (“uniform”) consisted of five uniform recombination maps of 10 Mb with recombination rate of 0.1, 0.25, 1, 4, and 10 times the mutation rate Sup. Fig. 5B. For the first set, we simulated 10 replicates of 150 diploid individuals (50 individuals per population). For the second and third sets, we simulated one replicate. We recorded the true ARGs in TreeSeq format, and also recorded haplotype data in VCF format using tskit version 0.4.1 (Kelleher et al., 2018).

### Demography inference

#### MSMC2

For the narrow high-rec. and uniform scenarios, we used four diploid individuals from each population for demography inference with MSMC2 (Malaspinas et al., 2016; Wang et al., 2020). For the stepwise scenario, we treated ten simulations as ten independent chromosomes, and downsampled four diploid individuals (eight haploid sequences) per population without replacement ten times as ten “replicates” (Note that they are not true independent replicates because they were sampled from a common ARG for each chromosome). Input multihetsep files were generated from the VCF file and masks for each chromosome of each downsample of each replicate using genrate_multihetsep.py of msmc-tools (Schiffels & Wang, 2020). We ran MSMC2 for each population or population pair to infer historical coalescence rates. The estimates of historical effective population size were obtained as the inverse of the inferred coalescence rate for each population, scaled with the true mutation rate of 4.6 *×* 10*^−^*^9^. The rCCR was obtained by dividing the between-populations coalescence rate with the average within-population coalescence rate. For visualization in Figs. 3, 4, we computed mean and standard deviation of the inferred effective population size and rCCR with a custom script.

#### Relate

For the stepwise scenario, we treated ten simulations as ten independent replicates. We applied filtering of variable sites based on the position according to masking conditions using BCFTools version 1.9 (Danecek et al., 2021). We inferred ARGs from the masked VCF using Relate version 1.1.6 (Speidel et al., 2019) specifying the true mutation rate, true recombination maps, and haploid population size of 2,000,000, and inferred demography with two iterations. The estimates of historical effective population size were obtained as the inverse of the inferred coalescence rate for each population, scaled with the true mutation rate of 4.6 *×* 10*^−^*^9^. The rCCR was obtained by dividing the coalescence rate between populations with the average within-population coalescence rate. For visualization in Figs. 3, 4, we computed mean and standard deviation of the inferred effective population size and rCCR with a custom script.

#### Stairway plot 2

We ran Stairway plot 2 version 2.1 (Liu & Fu, 2020) for the stepwise scenario. We treated ten simulations as ten independent replicates. We split the VCF by population applying masks with VCFTools version 0.1.16 (Danecek et al., 2011). We computed the unfolded SFS and prepared blueprint configuration files using custom scripts, and ran Stairway plot 2 with default parameter values. For visualization in Fig. 3, we computed mean and standard deviation of the inferred effective population size with a custom script.

### Coalescence time analysis

#### MSMC2

We focused on two haploids of the first chromosome (simulation run) of the first down- sample in pop1, and compared true TMRCA recorded in the true ARG (in TreeSeq format) and inference by MSMC2. We extracted the true TMRCA of the focal pair of haploid genomes in TreeSeq with tskit. To obtain inference by MSMC2, we ran the decode program of MSMC2 with decode -m 0.0092 -r 0.00736 -I 0,1 -t 32 -s 1000. Based on the output of decode, we recorded the index of epoch with the highest probability for each window. We aligned true and inferred TMRCA treating an intersected range as a unit, and computed Spearman’s correlation coefficient in R version 4.3.1 (R Core Team, 2023).

#### Relate

We focused on TMRCA of the entire genealogy of 300 haploid genomes of the first simulation replicate. We extracted TMRCA along the chromosome from the true ARG in TreeSeq using tskit. To obtain TMRCA along the chromosome of the ARG inferred by Relate, we converted the genealogies (in mut and anc format) to TreeSeq using RelateFileFormats program in Relate, and extracted TMRCA along the chromosome using tskit. We aligned true and inferred TMRCA treating an intersected range as a unit, and computed Spearman’s correlation coefficient in R.

### Empirical study

#### Data

We used phased whole-genome resequencing (WGR) data of 179 blackcaps (Ishigohoka et al., 2023), and unphased five garden warblers and three African hill babblers (Delmore et al., 2020). We computed mean recombination rate in 10-kb sliding windows along the blackcap genome based on (Bascón-Cardozo et al., 2024).

#### Demography inference MSMC2

We first applied callability masks to the blackcap genome and defined high- and low- recombining halves of the genome for each population or population pair. Specifically, we chose for each population at most four individuals with mean read depth of at least 15x, excluding pairs of related individuals based on kinship coefficient (Manichaikul et al., 2010) computed using relatedness2 option in VCFTools. We created a mask file per individual using bamCaller.py of msmc-tools (Schiffels & Wang, 2020) and merged them for each population or population pair using bedtools merge (Quinlan & Hall, 2010). The mask for each population or population pair was applied on the blackcap recombination map (Bascón- Cardozo et al., 2024), and we ordered genomic intervals within the unmasked regions according to the recombination rate. Regions in the first and the second halves were defined as the lower- and higher-recombining halves.

After defining the regions to be used for inference, input multihetsep files were generated from the phased VCF and the mask file using genrate_multihetsep.py of msmc-tools (Schiffels & Wang, 2020). We ran MSMC2 for each population or population pair to infer historical coalescence rates. The estimates of historical effective population size were obtained as the inverse of the inferred coalescence rate for each population, scaled with a mutation rate of 4.6 *×* 10*^−^*^9^ estimated in the collared flycatcher (Smeds et al., 2016). The rCCR was obtained by dividing the between-population coalescence rate with the average within-population coalescence rate.

#### Relate

Relate requires haplotype data with polarized mutations. We polarized biallelic SNPs in blackcaps using allele frequencies in two outgroup species, garden warblers (n=5) and African hill babblers (n=3). Specifically, after removing SNPs with more than two alleles including the three species, we split blackcap SNPs into the following five categories.

1. Sites at which all garden warblers had missing genotype.
2. Sites fixed in garden warblers
3. Sites segregated among garden warblers and missing in all African hill babblers
4. Sites segregated among both garden warblers and African hill babblers
5. Sites segregated among garden warblers and fixed in African hill babblers

For each category we applied the following heuristics to polarize mutations. For sites of the first type, we defined the minor allele among blackcaps to be the derived state (i.e. the major allele is the ancestral state). For sites of the second type, we defined the allele possessed by garden warbler to be the ancestral state. For sites of the third or fourth type, we defined the minor allele among blackcaps to be the derived state (i.e. the major allele is the ancestral state). For sites of the fifth type, we defined the allele possessed by African hill babbler to be the ancestral state.

We defined low- and high-recombining halves of the genome in BED format based on the blackcap recombination map (Bascón-Cardozo et al., 2024). Based on these BED files to mask high/low-recombining half and repeats retrieved from UCSC Genome Browser tracks (Raney et al., 2023) for the blackcap assembly (GenBank: GCA_009819655.1), we made a mask file in FASTA format for each condition using BEDTools maskfasta. Using the phased and polarized VCF, recombination map, and the mask, we ran Relate to infer genealogies with mutation rate of 4.6 *×* 10*^−^*^9^ and effective population size of 500,000. We inferred demography from the genealogies using RelateCoalescenceRate program of Relate with mutation rate of 4.6 *×* 10*^−^*^9^ and five times of iterations. The estimates of historical effective population size were obtained as the inverse of the inferred coalescence rate for each population with rescaling of time by generation time of 2 years (Delmore et al., 2020). The rCCR was obtained by dividing the between-population coalescence rate with the average within-population coalescence rate.

## Acknowledgments

This work was supported by the Max Planck Society (Max Planck Research Group grant MFFALIMN0001 to ML), and the DFG (project Nav05 within SFB 1372 – Magnetoreception and Navigation in Vertebrates (395940726) to ML). We thank Julien Dutheil, Linda Odenthal-Hesse, and Diethard Tautz for feedback.

## Data availability

Scripts used for the simulations and input data, processed output data and scripts for the empirical analyses are found in Zenodo (https://doi.org/10.5281/zenodo.10613446).

